# A flagellar accessory protein links chemotaxis to surface sensing

**DOI:** 10.1101/2024.06.20.599946

**Authors:** Rachel I. Salemi, Ana K. Cruz, David M. Hershey

## Abstract

Bacteria find suitable locations for colonization by sensing and responding to surfaces. Complex signaling repertoires control surface colonization, and surface contact sensing by the flagellum plays a central role in activating colonization programs. *Caulobacter crescentus* adheres to surfaces using a polysaccharide adhesin called the holdfast. In *C. crescentus*, disruption of the flagellum through interactions with a surface or mutation of flagellar genes increases holdfast production. Our group previously identified several *C. crescentus* genes involved in flagellar surface sensing. One of these, called *fssF*, codes for a protein with homology to the flagellar C-ring protein FliN. We show here that a fluorescently tagged FssF protein localizes to the flagellated pole of the cell and requires all components of the flagellar C-ring for proper localization, supporting the model that FssF associates with the C-ring. Deleting *fssF* results in a severe motility defect that we show is due to a disruption of chemotaxis. Epistasis experiments demonstrate that *fssF* promotes adhesion through a stator-dependent pathway when late-stage flagellar mutants are disrupted. Separately, we find that disruption of chemotaxis through deletion of *fssF* or other chemotaxis genes results in a hyperadhesion phenotype. Key genes in the surface sensing network (*pleD*, *motB*, and *dgcB*) contribute to both Δ*flgH-*dependent and Δ*fssF-* dependent hyperadhesion, but these genes affect adhesion differently in the two hyperadhesive backgrounds. Our results support a model in which the stator subunits of the flagella incorporate both mechanical and chemical signals to regulate adhesion.

**Importance:** Biofilms pose a threat in clinical and industrial settings. Surface sensing is an early step in biofilm formation. Studying surface sensing can help develop strategies for combating harmful biofilms. Here, we use the freshwater bacterium *Caulobacter crescentus* to study surface sensing. We characterize a previously unstudied gene, *fssF*, and find that it localizes to the cell pole in the presence of three proteins that make up a component of the flagellum called the C-ring. Additionally, we find that *fssF* is required for chemotaxis but dispensable for swimming motility. Lastly, our results show that mutating *fssF* and other genes required for chemotaxis causes a hyperadhesive phenotype. We propose that surface sensing requires chemotaxis for a robust response to a surface.

## Introduction

Bacteria frequently encounter solid surfaces in their surroundings that offer favorable habitats for colonization (Dang and Lovell, 2016; Harshey and Partridge, 2015; O’Toole and Wong, 2016). Attachment to surfaces provides improved access to nutrient resources and protection from environmental stressors. These benefits are vital for growth and fitness in many bacteria. As bacteria undergo development, they also transition from a motile state to a sessile, attached state (O’Toole et al., 2000). The attached lifestyle poses a significant threat in clinical and industrial settings (Muhammad et al., 2020). Understanding the mechanisms underlying surface colonization can inform new strategies to control bacterial colonization.

The motile to sessile transition requires signaling pathways that respond to physical contact with surfaces. Many bacteria use a molecular machine called the flagellum to recognize surface contact (Belas, 2014; Berne et al., 2018; Haiko and Westerlund-Wikström, 2013) (Figure 1). This sophisticated trans-envelope complex is best known for its role in cellular motility. The flagellar components are divided into four classes based on an assembly hierarchy. Class I genes encode for master regulators that control transcription of early flagellar components. Class II genes encode the cytoplasmic membrane associated motor proteins, class III genes encode the type III secretion system (T3SS) and hook basal body (HBB), and class IV genes encode flagellin proteins that make up the filament. Each class of flagellar genes requires proper expression and assembly of the previous class in the hierarchy (Aldridge, 2002; Belas, 2014; Minamino and Imada, 2015; Wu and Newton, 1997) (Figure 1B). The flagellar motor contains a cytoplasmic-ring (C-ring) composed of FliG, FliM, and FliN and a membrane-spanning ring (MS-ring) composed of FliF that form the rotor complex (Belas, 2014; Henderson et al., 2020; Khan et al., 1992; Kubori et al., 1997; Minamino and Imada, 2015; Zhao et al., 1996). Ion channels called stators engage with the C-ring to turn the rotor and its associated filament (Chang et al., 2020; Sridhar, 2020). Stator subcomplexes are composed of MotA and MotB, and they can rotate the motor in the in one direction to propel a cell forward or in the the opposite direction to cause the cell to reorient (Chang et al., 2020; Koyasu and Shirakihara, 1984; Liu et al., 2014).

**Figure 1:**
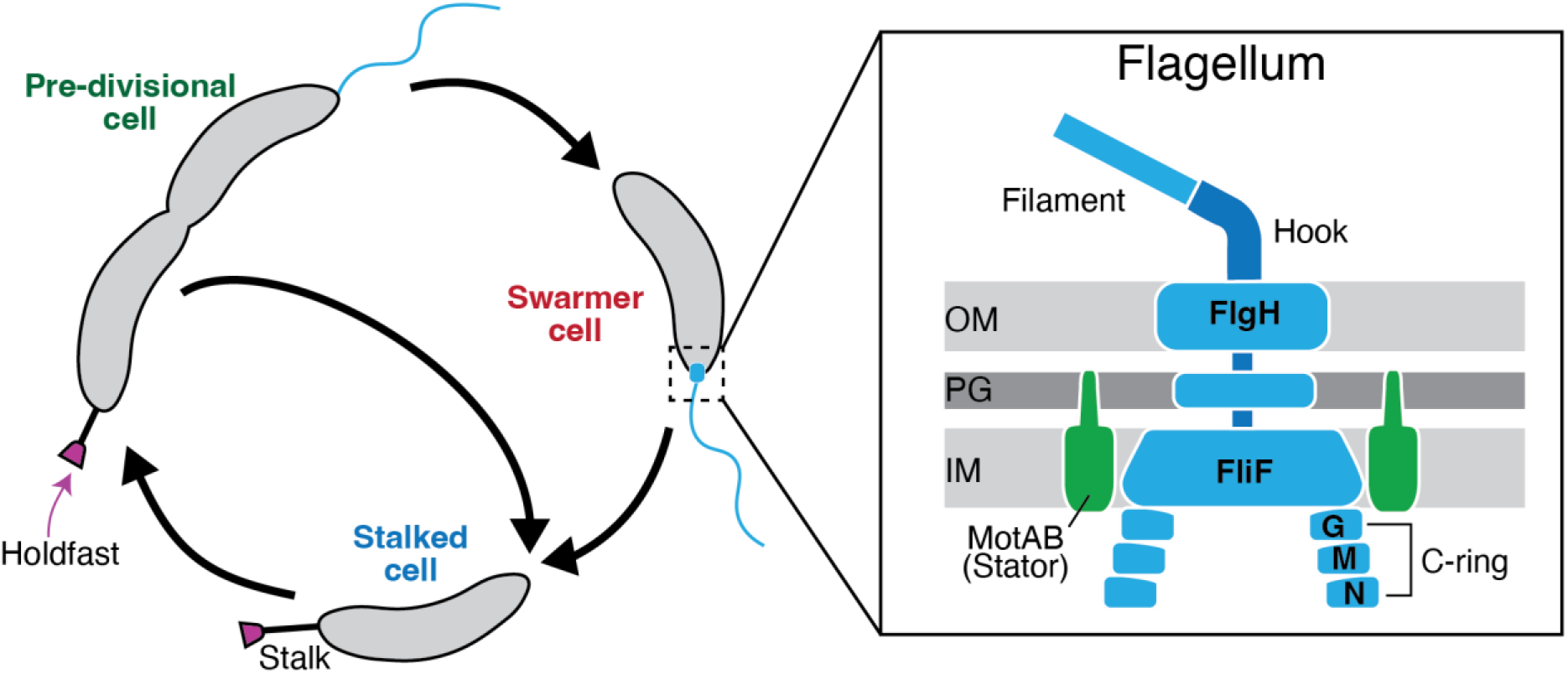
The *Caulobacter crescentus* flagellum is a complex multimeric machine. *C. crescentus* cells begin as a motile swarmer cell displaying a flagellum but are unable to reproduce. Differentiation into a stalked cell allows the cell cycle to continue and replication to begin. During this differentiation, the flagellum is lost and replaced by a stalk. Stalked cells can also produce an adhesive polysaccharide called the holdfast which allows them to irreversibly attach to a surface. Stalked cells divide to release a swarmer cell into the environment. The flagellum is composed of many subunits and spans the entire cell envelope. The cytoplasmic C-Ring is composed of three proteins that bind in a hierarchical manner, FliG (G), FliM (M), and FliN (N). The MS-ring spans the inner membrane and is comprised of the protein FliF. The L-Ring spanning the outer membrane is comprised of the protein FlgH.

The direction of motor rotation is regulated by a behavior called chemotaxis. Chemotaxis allows cells to move towards attractants or away from repellents (Chang et al., 2020). The *E. coli* chemotaxis system utilizes receptors called methyl-accepting chemotaxis proteins (MCPs) that detect attractants in the environment (Bi and Sourjik, 2018; Hansen et al., 2008; Huang et al., 2019; Szurmant and Ordal, 2004). Attractant binding controls a histidine kinase, CheA, that phosphorylates the response regulator protein CheY (Hansen et al., 2008; Muok et al., 2020). CheY-P translocates to the flagellar motor and interacts with the C-ring proteins FliM and FliN (Haiko and Westerlund-Wikström, 2013; Sarkar et al., 2010). This interaction disrupts the smooth swimming pattern by switching the direction of motor rotation, causing the cells to reorient (Koyasu and Shirakihara, 1984; Liu et al., 2014). Though the *E. coli* system has proven instrumental in determining the general principles of chemotaxis, bacteria such as *Caulobacter crescentus* harbor expanded chemotaxis systems with many copies of these core proteins (Berne and Brun, 2019; Nesper et al., 2017)

The rotation of the flagellum becomes physically obstructed during surface encounters (McCarter et al., 1988). This blocked state activates signaling pathways that promote surface adaptation by altering motility behavior, increasing irreversible attachment, and promoting biofilm formation (Belas, 2014; Berne et al., 2018; Hershey et al., 2021). The second messenger cyclic di-guanosine monophosphate (c-di-GMP) plays a central role in promoting surface adaptation (Romling *et al*. 2013, Jenal *et al*. 2017, Valentini and Filloux 2016). High concentrations of c-di-GMP promote biofilm formation and sessile behaviors, while low concentrations are associated with increased motility. Cellular concentrations of c-di-GMP are regulated by two enzymic activities: Diguanylate cyclase (DGCs) and Phosphodiesterase (PDEs) which respectively synthesize and degrade c-di-GMP (Ross et al., 1987). Though the enzymatic activity of DGCs is well studied, the flagellum regulates c-di-GMP production through complex signaling pathways that have been difficult to disentangle (Hershey, 2021).

*Caulobacter crescentus* is a robust model organism for studying surface colonization (Berne et al., 2018; Bodenmiller et al., 2004). This gram-negative, alpha-proteobacterium is abundant in freshwater systems (Bodenmiller et al., 2004). During the *C. crescentus* cell cycle, sessile cells called stalked cells divide asymmetrically to release non-replicative swarmer cells (Figure 1) (Schniederberend et al., 2019). Swarmer cells harbor a flagellum and pili at the old cell pole that are used for motility and surface sensing (Ellison et al., 2017; Hug et al., 2017; Skerker and Laub, 2004). Swarmer cells must differentiate into stalked cells before they can replicate (Jenal, 2000). Swarmer cells transition to become stalked cells by releasing their flagella, retracting their pili, synthesizing a stalk, and initiating chromosome replication (Skerker and Laub, 2004). During the swarmer-to-stalked transition *C. crescentus* cells can synthesize a specialized adhesin called the holdfast at the stalked cell pole (Berne et al., 2018; Chepkwony and Brun, 2021; Li et al., 2012). Surface sensing stimulates adhesion by increasing the likelihood that stalked cells will assemble a holdfast, making holdfast production an ideal phenotype for dissecting surface sensing (Hershey et al., 2021; Li et al., 2012).

We previously identified a large set of *C. crescentus* genes involved in signaling pathways activated by the flagellum (Hershey et al., 2021). Mutations in flagellar assembly genes stimulate the surface sensing pathway, indicating that disrupting the flagellar motor through mutation can mimic mechanical cues for surface contact. The resulting gain of function phenotype was used to identify *flagellar <u>s</u>ignaling <u>s</u>uppressor* (*fss*) mutations that restore wild-type levels of adhesion in a hyper-adhesive flagellar mutant background (Δ*flgH*). These novel surface sensing genes make up two, separate pathways. A “developmental” pathway requiring the DGC *pleD* is activated when flagellar assembly is disrupted at any stage. A second, “mechanical” pathway requiring the MotAB stators and the stator-associated DGC *dgcB* is activated specifically in late-stage flagellar mutants that retain an intact motor complex. Deletion of genes required for early stages of flagellar assembly (i.e. *fliF*, *fliG*, *fliN*, *fliM*) results in a hyperadhesivie phenotype that is entirely dependent developmental pathway. Deletion of genes required for assembly of the hook and filament (i.e *flgH*) results in a hyperadhesive phenotype that involves both the developmental and mechanical pathways.

Here we have examined a surface sensing factor called *fssF* (CC_2175, CCNA_02257) that contributes to holdfast stimulation downstream of the Δ*flgH* mutation. FssF shows high sequence homology to the C-ring protein FliN, and we find that FssF localizes to the flagellated pole of *C. crescentus* cells in a C-ring dependent manner. Mutating *fssF* abolishes the ability of cells to migrate through soft agar and eliminates chemotaxis behavior in liquid. When adhesion is activated by deletion of *flgH,* we find that *fssF* contributes to the stator-dependent, mechanical pathway. Separately, we find that Δ*fssF* and other mutants that cannot perform chemotaxis exhibit a hyperadhesive phenotype that is distinct from that of flagellar mutants. Hyperadhesion in Che-mutants is entirely dependent on *motB* for activation. This work expands on understanding of bacterial surface responses by indicating that the flagellar motor integrates mechanical and chemical cues to promote surface colonization.

## Results

### FssF shows homology to the C-ring protein FliN

Deleting *flgH* or other flagellar genes results in a hyperadhesive phenotype in *C. crescentus* (Hershey et al., 2019). To identify genes required for this hyperadhesion phenotype, we previously performed adhesion profiling of a Δ*flgH* transposon library (Hershey et al. 2021). One of the genes identified through this screen was an uncharacterized gene that we named *fssF* (*flagellar <u>s</u>ignaling <u>s</u>uppressor F*) (Figure S1A).

*fssF* codes for a predicted 103 amino acid protein. To further understand the function of FssF, we compared protein sequences of other annotated flagellar proteins and found that the protein shows homology to FliN proteins from many bacteria, suggesting that *C. crescentus* contains two FliN homologs (Figure S1B). While the core C-ring proteins are conserved among flagellated bacteria, some C-ring complexes contain accessory proteins that fulfill specialized roles (Bischoff and Ordal, 1992; Henderson et al., 2020). We hypothesized due to the similarity with FliN that FssF may localize to the C-Ring and play a role in motility.

### C-Ring proteins are required for FssF-Venus localization

The C-ring complex in most bacteria is composed of three proteins: FliG, FliM, and FliN (Chen et al., 2011)(Figure 1). In *E. coli*, C-ring proteins bind in a hierarchical manner, meaning that FliG must assemble first to create a scaffold for FliM and FliN to bind (Kubori et al., 1997). Thus, deletion of FliG would be predicted to cause mislocalization of all downstream C-ring components (Figure 2A). We confirmed this model in *C. cresentus* by fluorescently labeling all three C-ring proteins (Figure 2B, S3). We generated C-terminally tagged forms of each C-ring gene by fusing *venus* to the 3’ end of each gene at its native locus. The tagged proteins supported wild-type levels of migration through soft agar, and western blotting showed that the tagged proteins accumulated in their full-length forms (Figure S2). We observed three types of localization patterns: polar localization, diffuse localization, and mislocalization. Strains that exhibited polar localization contained foci specifically at the cell pole in approximately 10-20% of cells. Strains that exhibited diffuse localization lacked foci. Strains that exhibited mislocalization contained many foci with no apparent preference for the poles. All three C-ring proteins exhibited polar localization in the wild-type background (Figure 2B, Figure S3). In a Δ*fliG* background, both FliM-Venus and FliN-venus displayed a mislocalization pattern (Figure S3). In a Δ*fliM* background, FliG-Venus localized to the cell pole, while FliN-Venus was diffuse (Figure S3). In a Δ*fliN* background, FliG-Venus exhibited polar localization while FliM-Venus exhibited a diffuse pattern (Figure S3). These results are consistent with previous literature that indicate FliG is required for localization of downstream C-ring proteins and that FliN and FliM associate with one another for assembly (Kubori et al., 1997).

**Figure 2:**
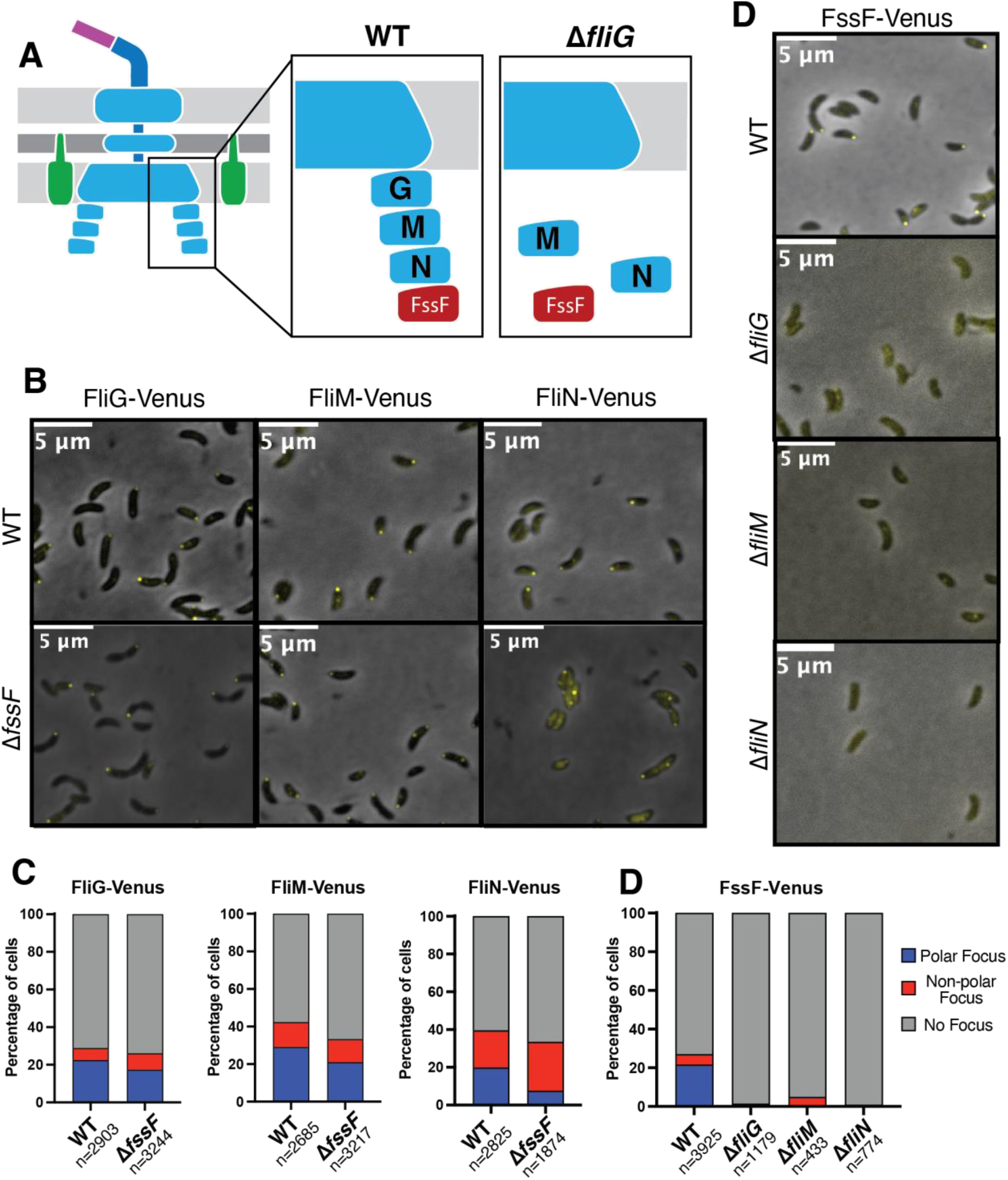
FssF requires the C-Ring for proper localization. (A) Model of hierarchical assembly of the C-Ring. FliG (G) must bind first, followed by FliM (M) and FliN (N). In the absence of FliG, localization of other C-ring proteins is lost. (B) Representative micrographs showing the subcellular localization of FliG-Venus, FliM-Venus, and FliN-Venus in a wild-type (WT) and Δ*fssF* background. (C) Quantification of localization of FliG-Venus, FliM-Venus, and FliN-Venus in a wild type background and Δ*fssF* backround. Cells were binned into three categories based on localization: Polar focus, non-polar focus, and no focus. (D) Representative micrographs showing the subcellular localization pattern of FssF-Venus in a wild type (WT), Δ*fliG*, Δ*fliM,* and Δ*fliN* backgrounds. (E) Quantification of localization of FssF-Venus in a wild type, Δ*fliG*, Δ*fliM*, and Δ*fliN* background.

To determine whether FssF localizes to the flagellum, we fluorescently tagged *fssF* at its native locus with *venus* and confirmed the functionality of this fusion using soft agar motility assays and western blots (Figure S2). We imaged this fusion in the wild-type, Δ*fliG*, Δ*fliM*, and Δ*fliN* backgrounds (Figure 2D-E). FssF-Venus localized to the flagellar pole in wild-type cells that retain all three canonical C-ring proteins and displayed a diffuse localization pattern when *fliG, fliM,* or *fliN* was deleted (Figure 2D-E). We found that in a wild-type background, 22% of cells contained a polar FssF-Venus focus, while 0% of cells contained a polar focus in any of the three C-Ring deletion backgrounds (Figure 2E). These results indicate that FssF requires a properly assembled C-ring for recruitment to the flagellar motor. Deletion of *fssF* did not affect the localization of FliG-Venus or FliM-Venus but increased the percentage of mislocalized FliN-Venus foci suggesting that *fssF* may affect FliN localization (Figure 2C).

### FssF is a dispensable for motility but required for chemotaxis

To understand the function of *fssF*, we examined motility phenotypes of the Δ*fssF* mutant using soft agar assays and live cell imaging. Δ*fssF* cells were unable to spread through semi-solid agar (Figure 3). Spreading in soft agar requires flagellar assembly, motor rotation and chemotaxis (Wolfe and Berg, 1989). To determine which of these processes was disrupted in the Δ*fssF* mutant, we observed cellular motility in liquid medium using live cell imaging. We compared the swimming phenotype in Δ*fssF* to the phenotypes of three other strains: Wild-type, Δ*flgH,* and Δ*cheYII.* Wild-type *C. crescentus* cells swim and change directions in previously described a run-reverse-flick sequence (Koyasu and Shirakihara, 1984; Liu et al., 2014), while Δ*flgH* cells are non-motile. Δ*cheYII* cells swim in liquid but do not display run-reverse-flick directional switching and often display spiraling trajectories indicative of being trapped in the hydrodynamic boundary (Conrad, 2012). We found that the swimming behavior of Δ*fssF* was indistinguishable from the Δ*cheYII* mutant as these cells are capable of swimming but lack directional switching (Figure 3). We conclude that the Δ*fssF* mutant does not migrate in soft agar due to an inability to perform chemotaxis.

**Figure 3:**
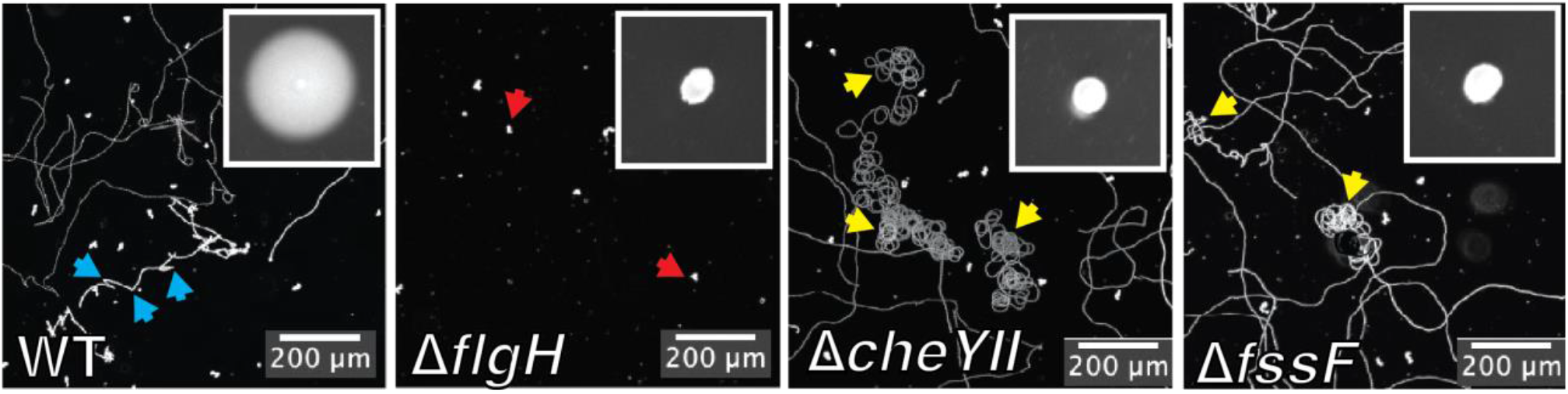
FssF is required for chemotaxis. FssF phenocopies a chemotaxis null mutant in swimming behavior. Maximum projections from live cell tracking of each strain. Immobile cells appear as white spots (Red arrows) and swimming trajectories appear as white lines. Directional switching is indicated by blue arrow, while cells unable to change direction are labeled by the yellow arrow. A representative soft agar motility assay is shown in the top right corner of each panel.

### *fssF* contributes to the stator branch of the *C. crescentus* surface sensing pathway

We previously determined that late-stage flagellar mutations such as Δ*flgH* activate adhesion by mimicking surface contact and stimulating two distinct signaling pathways (Figure 4A) (Hershey et al., 2021). To determine if deletion of *fssF* disrupts the developmental or the mechanical pathway, we deleted *fssF* in an early-stage (Δ*fliF*) and a late-stage (Δ*flgH*) flagellar mutant background. If *fssF* were involved in the developmental pathway, we would expect deletion of *fssF* to suppress hyperadhesion in both the Δ*flgH* and Δ*fliF* backgrounds. If *fssF* were involved in the mechanical pathway, we would expect Δ*fssF* to suppress only Δ*flgH*-mediated hyperadhesion. We used crystal violet (CV) staining to perform these epistasis experiments (Figure 4B). There was no difference in adhesion between the Δ*fliF* mutant and the double Δ*fliF* Δ*fssF* mutant. However, hyperadhesion in the Δ*flgH* strain was significantly reduced by the deletion of *fssF* (Figure 4B). These results indicate that *fssF* controls adhesion by supporting the *motB*-dependent, mechanical pathway.

**Figure 4:**
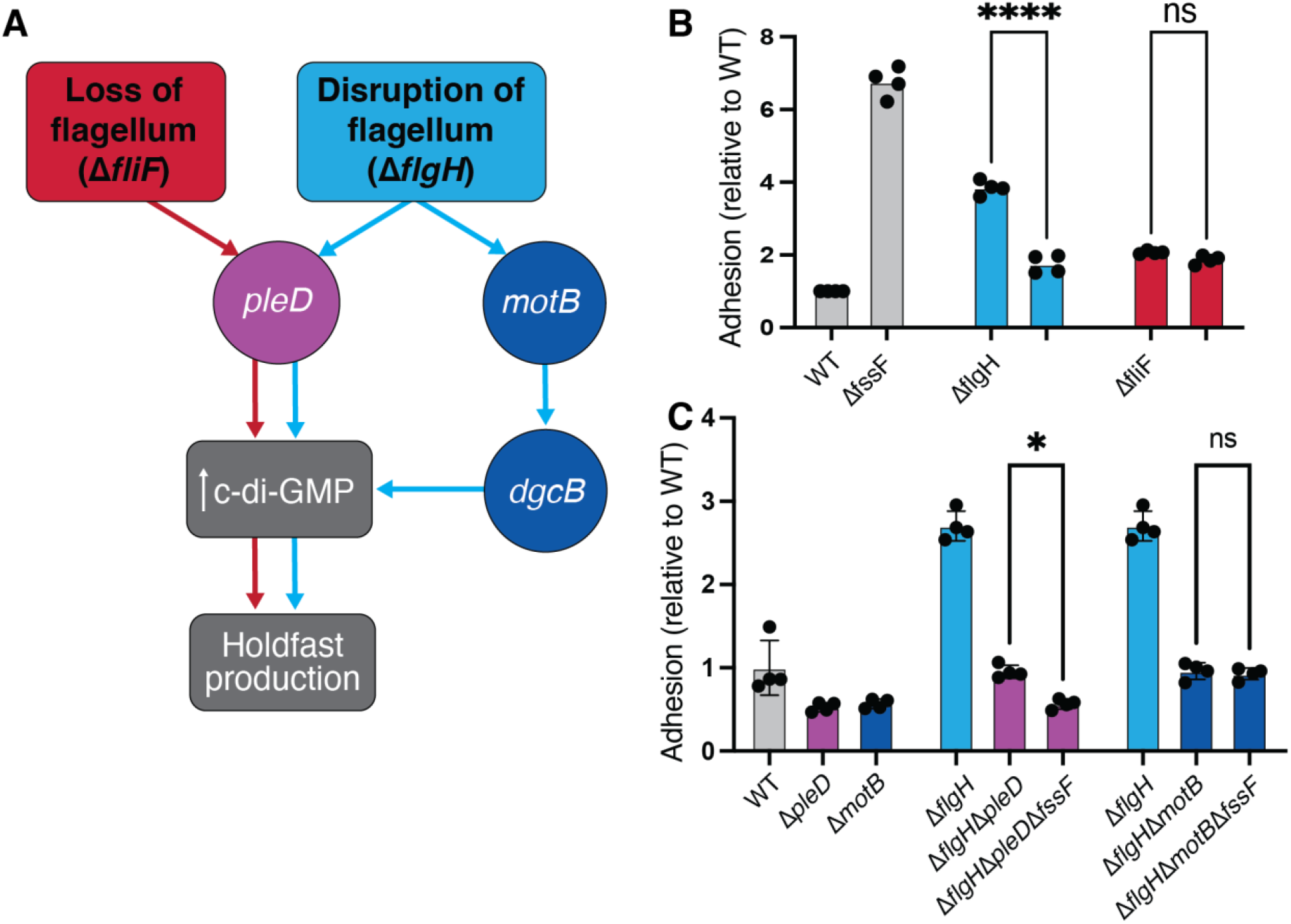
*fssF* is required for full activation of the *motB-*dependent adhesion pathway. (A) Model of hyperadhesion regulation induced by loss or disruption of flagellar machinery. When the flagellum is lost due to mutations in class II flagellar genes such as *fliF* or the C-ring genes, the *pleD*-dependent adhesion pathway is stimulated. Deletions in genes in the *motB*-dependent adhesion pathway do not affect hyperadhesion in class II flagellar gene mutations. The deletion of class III and IV flagellar genes such as *flgH* result in a hyperadhesion phenotype that is dependent on both *pleD* and *motB*. (B) and (C) CV staining to measure adhesion relative to wild type. Statistical significance determined by one-way ANOVA test and Tukey’s multiple-comparison test. The results demonstrate that *fssF* supports Δ*flgH-*dependent hyperadhesion through the mechanical pathway.

To confirm that *fssF* is required for signaling through the mechanical pathway, we deleted *fssF* in a Δ*flgH* Δ*pleD* background and a Δ*flgH* Δ*motB* background. We expected the Δ*pleD* and Δ*fssF* mutations to have additive effects in suppressing hyperadhesion in the Δ*flgH* background, confirming that these two genes promote hyperedhesion through distinct pathways. Indeed, a Δ*fssF* Δ*flgH* Δ*pleD* triple mutant displayed lower adhesion than the Δ*flgH* Δ*pleD* strain. Deletion of *fssF* did not affect adhesion in the Δ*flgH*Δ*motB* background, indicating that *fssF* and *motB* promote adhesion through the same pathway in the Δ*flgH* background (Figure 4C).

### Disruption of chemotaxis produces a novel hyperadhesive phenotype

While performing the epistasis experiments described above, we observed that the Δ*fssF* mutation stimulated adhesion in the wild-type background (Figure 4B). This phenotype is unexpected for suppressors of Δ*flgH* hyperadhesion. These genes are thought to support elevated adhesion when surface-sensing is activated, and deleting these genes in the wild-type background typically either does not affect or lowers adhesion(Hellenbrand et al., 2024; Hershey et al., 2021). We predicted that the Δ*fssF* strain showed elevated adhesion because disrupting chemotaxis increases holdfast production (Figure 5SA). Because both Δ*fssF* and Δ*cheYII* show a Che-motility phenotype, we compared adhesion in the Δ*fssF*, Δ*cheYII* and double Δ*fssF* Δ*cheYII* mutants using CV staining and direct microscopic quantification of holdfast production (Figure 5A and FigS4). All three strains exhibited identical hyperadhesive phenotypes, demonstrating that abolishing the ability of *C. crescentus* cells to perform chemotaxis increases adhesion. To ensure that this phenotype was due to disruption of chemotaxis and not specific to Δ*cheYII*, we also measured adhesion in Δ*cheAI* and found that this strain also displayed a hyperadhesion phenotype (Figure S4A). The Δ*fssF* hyperadhesion phenotype and motility defect could be complemented by expressing the gene from its native promoter at the xylose locus (Figure S4).

**Figure 5:**
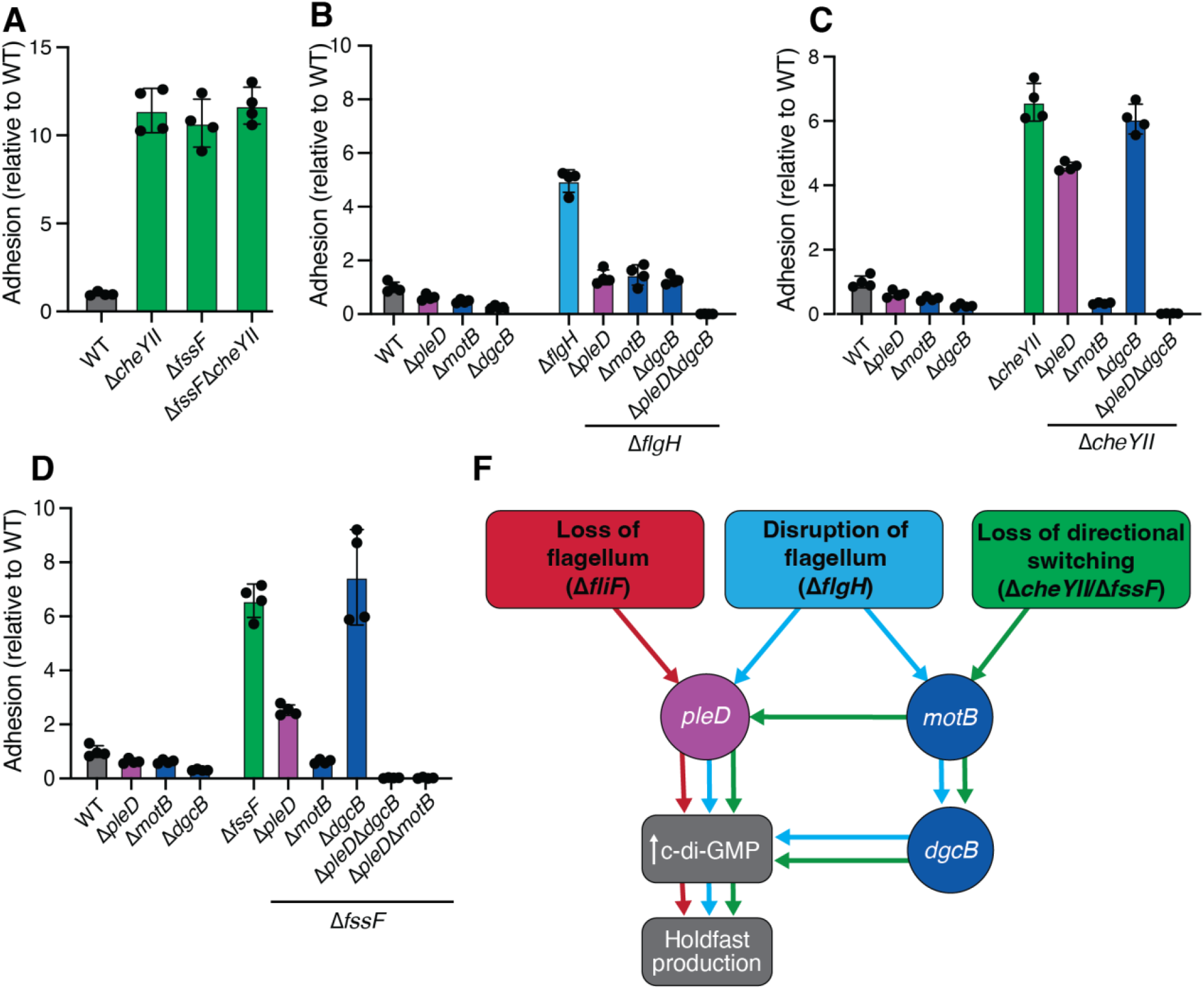
*ΔfssF* hyperadhesion differs from Δ*flgH* hyperadhesion. In panels A-D, CV staining to compare adhesion in various mutants to wild type. (A) Che-mutants show a hyperadhesive phenotype. (B) Deleting *pleD, motB* or *dgcB* in the Δ*flgH* mutant background modestly suppresses the hyperadhesvie phenotype. A Δ*flgH* Δ*pleD* Δ*dgcB* triple mutant is nonadhesive, showing that *dgcB* and *pleD* play separate, additive roles in activating adhesion when the flagellar filament is disrupted. (C and D) In Che-backgrounds (Δ*fssF* and Δ*cheYII* are shown), deleting *pleD* provides minimal suppression of the hyperadhesive phenotype. *motB* is required for adhesion in Che-mutants, while *dgcB* is dispensable. Δ*cheYII* Δ*pleD* Δ*dgcB* and Δ*fssF* Δ*pleD* Δ*dgcB* triple mutants are nonadhesive, showing that *dgcB* and *pleD* play redundant roles in activating adhesion when chemotaxis is disrupted. (E) Updated model of adhesion regulation incorporating chemotaxis null hyperadhesion. In che-strains, we find that *pleD* and motB play an important role, and that *motB* activates both *pleD* and *dgcB*.

We sought to understand if regulators of Δ*flgH* hyperadhesion (*pleD, motB,* and *dgcB*) are also involved in the Δ*fssF*/Δ*cheYII* hyperadhesion and performed additional epistatsis experiments. As previously described, both *pleD* and *motB* contribute to adhesion in the Δ*flgH* (Hershey et al., 2021). The effects of the Δ*dgcB* mutation were identical to Δ*motB* in the Δ*flgH* background. A Δ*flgH* Δ*pleD* Δ*dgcB* triple mutant is non-adhesive (Figure 5B), showing that *pleD* and *dgcB* promote adhesion through separate, additive pathways in the Δ*flgH* background.

Deletion of *motB* in the Δ*fssF* and Δ*cheYII* backgrounds nearly abolished adhesion, while deletion of *dgcB* had no effect (Figure 5C & D). Deletion of *pleD* weakly suppresses hyperadhesion in Δ*cheYII* and Δ*fssF*. Though *dgcB* and *pleD* deletions do not show strong suppression of hyperadhesion individually, deleting both genes simultaneously in the Δ*cheYII*/Δ*fssF* backgrounds abolishes adhesion entirely. This indicates that *pleD* and *dgcB* play redundant functions in activating adhesion when chemotaxis is disrupted. We also measured adhesion of Δ*pleD*, Δ*dgcB*, and Δ*motB* in the Δ*cheAI* background and the results phenocopied our findings for Δ*cheYII* (Figure S4B-C). These results suggest that inducing hyperadhesion by disrupting chemotaxis activates a signal transduction sequence that is distinct from the one required for Δ*flgH* hyperadhesion, though the key genes involved remain the same.

## Discussion

Bacteria have robust mechanisms for colonizing solid surfaces, and flagella play crucial roles throughout the colonization sequence (Belas, 2014). Obstruction of the flagellar filament increases recruitment of the stators which is thought to trigger surface adaptation processes such as adhesion and biofilm formation (Baker and O’Toole, 2017; Lele et al., 2013; Li et al., 2012; McCarter et al., 1988; Tipping et al., 2013). Signal transduction pathways activated by obstruction of the flagellum stimulate the production of the second messenger c-di-GMP, but a more thorough understanding of how the downstream signaling networks are organized is needed.

In this study, we characterized a novel surface sensing protein called FssF from the freshwater bacterium *Caulobacter crescentus. fssF* was identified in a screen for signaling factors that activate holdfast production downstream of the flagellum (Hershey et al., 2021), and its predicted protein sequence shows high homology to the C-ring protein FliN. We found that fluorescently-tagged FssF localizes to the flagellar pole of *C. crescentus* cells in a manner dependent on all of the canonical C-Ring proteins (FliG, FliM, and FliN).

Additionally, Δ*fssF* cells retain the ability to swim in liquid but do not perform the directional switching that is characteristic of chemotaxis. Swimming behavior in Δ*fssF* cells is indistinguishable from an established chemotaxis mutant (Δ*cheYII*), confirming that FssF is required for chemotaxis.

The requirement of *fssF* for directional switching and the C-ring dependent localization of FssF-Venus to the cellular pole support the model that FssF is an accessory protein that associates with the C-ring of the flagellum. Diversifications and expansions of C-ring protein repertoires have been identified in other bacteria (Henderson et al., 2020). Our data suggest that duplication of the *fliN* gene led to subfunctionalization of FliN in *C*.

*crescentus* by separating the function of flagellar assembly from the function of directional switching. FliG, FliM, and FliN are required for assembly of the core C-ring structure, while FssF is specifically required for chemotaxis. We speculate that either FssF itself or a site on an FssF-dependent C-ring conformation serves as the binding site for the major CheY-like protein (CheYII) in *C*. *crescentus*.

Current models for surface sensing emphasize the flagellar motor’s ability to act as a mechanosensor by recruiting additional stator subunits as load on the filament increases (Lele et al., 2013; Tipping et al., 2013). Our previous work supported this model by showing that the stator genes are required for the increase in holdfast production when the surface sensing pathway is activated (Hershey et al., 2021). We also identified a second stator-independent pathway activated by disruption of the flagellum that required the developmental regulator *pleD* (Figure 4A). Here, we used epistasis analysis to show that *fssF* contributes specifically to the stator-dependent, mechanical branch of the *C. crescentus* surface-sensing network. Given that *fssF* is required for directional switching during chemotaxis (Figure 3), our results indicate that a robust surface response requires both load-dependent mechanosensing and directional switching.

We also identified a role for chemotaxis genes that is distinct from their function in the mechanical sensing pathway activated by deletion of *flgH*. Mutations disrupting chemotaxis (Δ*fssF,* Δ*cheYII,* and Δ*cheAI*) stimulate holdfast production in the wild-type background (Figure 5A). In contrast with our data, Berne and Brun reported that a Δ*cheAI* mutant shows a modest decrease in holdfast production (Berne and Brun, 2019). While we cannot explain the discrepancy in adhesion phenotypes of the two *C. crescentus* Δ*cheAI* mutants, we have confidence that our Che-mutants show the proper phenotypes. Each of our Che-mutants displays the expected non-spreading phenotype in soft agar (Figure 3) (Skerker et al., 2005; Wolfe and Berg, 1989), and another study recently reported a genetic screen that identified a hyperadhesive phenotype in a *cheAI* mutant (Zappa et al., 2023). Further, our dissection of the genetic basis for hyperadhesion and confirmation via counting holdfast (Figure S5) provides additional support for our assignment of holdfast phenotypes.

We used epistasis analysis to show that the Δ*flgH* and Δ*fssF* mutations activate holdfast production differently. In Che- (Δ*fssF,* Δ*cheYII,* and Δ*cheAI*) backgrounds, deleting *pleD* causes a slight reduction of the hyperadhesive phenotype and deleting *dgcB* has no effect. However, the Δ*fssF* Δ*pleD* Δ*dgcB* and Δ*cheYII* Δ*pleD* Δ*dgcB* triple mutants are completely non-adhesive. These results indicate that the two DGCs play redundant roles in activating adhesion when chemotaxis is disrupted. Further, we find that deletion of *motB* nearly abolishes adhesion in the Che-backgrounds. This data indicates that *motB* is upstream of both DGC enzymes. The relationships between these genes differs when hyperadhesion stimulated by deleting *flgH*. Deleting *motB, pleD,* or *dgcB* in the Δ*flgH* background reduces adhesion to the wild-type level. Δ*flgH* Δ*pleD* Δ*dcgB* and Δ*flgH* Δ*pleD* Δ*motB* triple mutants are completely nonadhesive, but the Δ*flgH* Δ*dgcB* Δ*motB* mutant retains wild type levels of adhesion. Thus, *pleD* and *dgcB* play separate, additive roles in activating adhesion when late stages of flagellar assembly are disrupted, and *motB* is upstream of *dgcB* but not *pleD.* The separation of the mechanical and developmental pathways is a key difference between adhesion regulation in Fla-mutants compared to Che-strains.

We propose that *pleD, motB, dgcB* and the downstream adhesion factors they control can be activated by multiple stimuli. Though all three genes appear central to a surface sensing network in *C. crescentus*, different sensory inputs seem to activate these genes in different ways. Disrupting the late-stages of flagellar assembly activates PleD and DgcB separately, causing these DGCs to perform separate, additive functions. Only DgcB is dependent on the stators in this context. Disrupting directional switching also activates both PleD and DgcB. However, these enzymes play redundant roles in activating adhesion and both appear dependent on the stators. We propose that the different epistasis patterns seen when holdfast production is activated by disrupting flagellar assembly or disrupting chemotaxis can be explained by network plasticity. The system appears to alter how signals are transduced in response to different stimuli.

Though we have not yet identified all genes involved in the surface sensing network, it is clear that the stators play a crucial role in directing flux through the system. Eliminating directional switching activates the stators to increase c-di-GMP production by stimulating *pleD* and *dgcB.* Disrupting late stages of flagellar assembly activates *dgcB* via *motB* and activates *pleD* through a separate mechanism (Hershey et al., 2021; Hug et al., 2017). Perhaps, ongoing debates surrounding the role(s) of stators in surface sensing can be explained by the fact that they signal differently in different contexts. Understanding how the stators direct flux through the surface sensing network will require characterizing additional genes required for stator-dependent (Che-) and stator-independent (Fla-) activation of *pleD*.

Our work emphasizes that the flagellar motor can integrate diverse sensory stimuli. Mechanical cues from the flagellar filament and chemical cues from chemotaxis systems are integrated into a single response. Recent studies have shown that mechanosensitive stator recruitment increases the affinity of activated CheY for the C-ring (Antani et al., 2021). Our analysis indicates that the connection between mechanosensing and directional switching may be more even sophisticated. Not only can load affect CheY binding to the C-ring, but CheY binding seems to also amplify stator-dependent responses to high load. These findings emphasize the role of the flagellum in environmental sensing. In addition to promoting cellular motility, this machine also serves as a sophisticated signaling hub that allows bacteria to process diverse sensory stimuli.

## Materials and Methods

### Bacterial growth and strain constructions

*C. crescentus* CB15 cells were grown in peptone yeast extract (PYE) broth shaking at 200 rpm overnight at 30°C unless otherwise indicated. Standard PCR and Gibson assembly (Gibson et al., 2009) were used to for cloning of deletion and insertion plasmids. *Escherichia coli* cultures were grown in Luria-Bertani (LB) broth supplemented with 50ug/mL kanamycin when required. Deletion and insertion plasmids were transformed into *C. crescentus* CB15 through electroporation followed by kanamycin selection (25ug/mL kanamycin) and a sacB-based counter selection using 3% wt/vol sucrose (Hmelo et al., 2015). Strains are listed in Table S1 and plasmids in Table S2. Primer sequences and details of plasmid construction are available upon request.

### Crystal Violet Staining Assay

*C. crescentus* cultures were grown overnight in PYE, then diluted to an optical density (OD660) of 0.5 with PYE. A 48-well plate was prepared with 450 ul of M2X media (1X M2 salts, 1% hunter base, 0.5 M CaCl2, 1M MgSO4, and 0.15% xylose) in each well (Hershey et al., 2019). Each well was inoculated with 1.5 ul of diluted culture. Each plate contained 4 replicates. Plates were sealed and grown at 30°C while shaking at 155 rpm for 17 hours. Cultures were then discarded, and the plates were washed continuously under running cold tap water for 1-2 min. Remaining adherent cells were stained by adding 500 ul of 0.01% crystal violet aqueous solution to each well and shaking the plates at room temperature for 10 min. The plates were washed for 1-2 min with running tap water. The remaining dye was dissolved by adding 500 ul of ethanol to each well and shaking for another 10 min. Staining was quantified by measuring the absorbance at 575 nm using a BioTek Synergy H1microplate reader. The absorbance value of each reading was first subtracted by the absorbance value of a blank sample. These values were then normalized and compared to the average WT absorbance value. Statistical analysis was performed through GraphPad Prism.

### Soft Agar Assay

*C. crescentus* PYE cultures were grown overnight and then diluted to an OD660 of 0.5 with PYE. 1.5 ul of diluted cultures were inoculated in PYE plates containing 0.3% agar. The plates were then left to incubate at 30°C for 72 hours. Images were taken using the Invitrogen iBright FL1500 Imaging System.

### Live Cell Microscopy to Analyze Swimming Behavior

2 ml PYE overnight cultures were backdiluted to an OD660 of 0.1. The cells were then grown to OD660 of 0.4-0.5, and diluted 1,000-fold in a microfuge tube containing 1 ml of PYE. Glass slides were inoculated with 2 ul of cells and enclosed with a coverslip and sealed with valap. Dark-field images were taken with a 10x objective lens every 50msec for 1 min in a Nikon Eclipse Ti series microscope and Orca Fusion BT digital CMOS camera (Hamamatsu). Maximum projections were calculated using Nikon NIS Elements software.

### Imaging Fluorescently tagged C-Ring proteins

Overnight PYE cultures of *C. crescentus* were backdiluted in M6HIGX (5 mM imidazole-HCl (pH 7.0), 2 mM sodium phosphate, 1% hunter base, 0.3% sodium glutamate-KOH (pH 7.0), and 0.3% xylose) to an OD660 of 0.1. At an OD660 of 0.4-0.5, 100 uL of culture was collected in a microcentrifuge tube and spun down for 1 minute at 3500 rcf. 85 uL of supernatant were removed and remaining pellet was resuspended by agitating the tube gently. 2 uL of the concentrated culture were plated on a 3% agarose pad. A Nikon Eclipse Ti series microscope with a 100x oil immersion objective lens was used to capture the fluorescently tagged proteins in cells. Fluorescence images were collected using a Prior Lumen 200 metal halide light source and a YFP-specific filter set (Chroma). Images were taken with 25-50 msec phase exposure and 3s fluorescence exposure. All images shown were processed using ImageJ and microbeJ was used for identification of cells and polar foci.

### Holdfast staining

*C. crescentus* cultures were grown overnight in PYE, then diluted 1:400 in M2X medium and grown overnight to an OD660 between 0.05-0.1. At the target OD, a sample of culture was taken and Alexa 594-WGA was added to a final concentration of 2ng/mL. The sample was then washed with 1mL of sterile H2O and centrifuged for 2 minutes at 6K xg. Supernatant was removed and cells were resuspended by agitating the tube gently. 2uL of sample were added to 3% agarose pad and imaged on Nikon Eclipse Ti series microscope with a 100x oil immersion objective lens with a 50 msec phase exposure followed by a 1-2 sec mCherry exposure. Fluorescence images were collected using a Prior Lumen 200 metal halide light source and a YFP- and mCherry-specific filter set (Chroma). All images shown were processed using ImageJ and microbeJ(Ducret et al., 2016).

### Immunoblotting

Cells were grown overnight in PYE and then backdiluted to and OD660 of 0.1. Cells from 0.5mL of culture (OD660 of 0.4-0.5) were collected by centrifugation. Pellets were resuspended in 1x TBS (20mM Tris, 0.15M NaCl, pH 7.5) with benzonase nuclease, incubated at 30°C for 10 minutes, and added to an equal concentration of 4x SDS-LOAD buffer containing 1%BME. Samples were boiled, loaded into 12% SDS-page gels, resolved and transferred to PVDF membrane via semi-dry transfer. Blocking was performed with 5% milk powder for 2 hours at room temperature. Membrane was then shaken overnight at 4°C in 5% milk containing 1:10000 dilution of anti-GFP serum from rabbit. The membrane was then subject to three 15 minutes washes in 1XTBST. Anti-rabbit HRP was applied in 5% milk at a concentration of 1:10000 and shaken at room temperature for 1 hr.

Gels were exposed on Invitrogen iBright FL1500 Imaging System using chemiluminescent substrate Western Lightning Plus by PerkinElmer.

## Acknowledgements

This work was supported by National Institutes of Health award R35GM150652 to D.M.H. and startup funds from the University of Wisconsin – Madison to D.M.H.

